# The Lon protease links nucleotide metabolism with proteotoxic stress

**DOI:** 10.1101/870733

**Authors:** Rilee D. Zeinert, Hamid Baniasadi, Benjamin Tu, Peter Chien

## Abstract

During stress all cells must maintain proteome quality while sustaining critical processes like DNA replication. In bacteria, the Lon protease is the central route for degradation of misfolded proteins. Here, we show that in *Caulobacter crescentus* Lon controls dNTP pools during stress through degradation of the transcription factor CcrM. We find that elevated dNTP/NTP ratios in Δ*lon* cells protects them from deletion of otherwise essential dTTP-producing pathways and shields them from lethality of hydroxyurea, known to catastrophically deplete dNTPs. Increased dNTP production in Δ*lon* results from higher expression of ribonucleotide reductase driven by increased CcrM. We show that misfolded proteins can stabilize CcrM by competing for limiting protease and Lon-dependent control of dNTPs improves fitness during protein misfolding conditions. We propose that linking dNTP production with the availability of Lon allows *Caulobacter* to maintain replication capacity when misfolded protein burden increases, such as during rapid growth or unanticipated proteotoxic stress.

**Highlights:**

- dCTP deaminase (DCD) is dispensable when Lon protease is absent due to increased dNTP pools.
- Stabilization of the Lon substrate CcrM transcriptionally upregulates ribonucleotide reductase, affording protection against hydroxyurea.
- Misfolded proteins can competitively inhibit CcrM degradation by the Lon protease.
- Titration of protein quality control is a mechanism that allows cells to respond to stresses that lack dedicated signal response pathways.

## Introduction

Protein quality-control (PQC) proteins such as chaperones and proteases are essential to protect cells from the harmful effects of protein misfolding or unfolding (Bukau et al., 2006; Sauer and Baker, 2011). The Lon protease is a conserved member of the bacteria PQC machinery that is responsible for ~ 50% of misfolded protein degradation (Goff et al., 1984). In order to degrade a large range of misfolded proteins, Lon is thought to recognize continuous regions of exposed hydrophobic sequences that are normally buried in folded proteins (Gur and Sauer, 2008), thus limiting the accumulation of potentially toxic misfolded proteins during stress conditions. During acute stresses, such as heat shock, the PQC system is deployed to limit damage and these stresses are often associated with a pause in cell division to provide opportunities for repair/response (Heinrich et al., 2016; Rowley et al., 1993). However, chronic PQC stresses that arise from increased misfolded protein burden such as that seen during rapid growth must be balanced with the need to support essential cell processes like DNA replication.

In addition to its role in PQC, Lon plays an important role during normal cell growth and division. In the α*-*proteobacterium *Caulobacter crescentus*, Lon degrades the replication initiator DnaA and the cell-cycle methyltransferase CcrM. Cells lacking Lon accumulate DnaA which results in excess chromosome content and the stabilization of CcrM results in aberrant transcription of cell cycle genes (Jonas et al., 2013; Wright et al., 1996). Lon also degrades the transcription factor SciP which represses expression of many cell cycle genes (Gora et al., 2013). In these cases, Lon recognizes specific sequence motifs as degrons because stabilizing mutations can be made without disrupting protein function or structure (Gora et al., 2013; Liu et al., 2019). Therefore, Lon can both degrade native substrates using specific degrons and destroy misfolded proteins by recognizing general features of poor quality substrates such as exposed hydrophobic regions (Mahmoud and Chien, 2018). Consideration of this large target repertoire is particularly important given substrate competition for a limited pool of protease during PQC stress conditions. It remains unknown whether there is an adaptive link in different functions of the Lon protease that may aid growth during chronic PQC stress conditions.

In this study, we show that a Lon-dependent circuit connecting PQC and nucleotide metabolism is important during PQC stress conditions. A transposon sequencing approach identified genes that are dispensable only in cells lacking Lon. One such gene is dCTP deaminase (DCD), which together with ribonucleotide reductase (RNR), is critical for dNTP production. We find that loss of DCD is tolerated in Δ*lon* strains because of elevated dTTP pools and metabolomics profiling showed higher ratios of dNTP/NTP in general. We found this change in the dNTP pool was due to increased RNR activity driven by the stabilization of the Lon substrate CcrM which transcriptionally upregulates RNR. This observation is confirmed by the increased tolerance of Δlon strains to hydroxyurea, a known inhibitor of RNR. Given the need for misfolded protein degradation during PQC stress, we considered that this Lon-dependent circuit may result in stabilization of CcrM when cells are faced with increased misfolded protein burden. We find that misfolded proteins can inhibit Lon degradation of CcrM *in vitro* and CcrM is stabilized during PQC stress *in vivo*, leading to increased RNR activity. Finally, we show that interrupting this circuit reduces fitness when the PQC system is taxed during rapid growth conditions, consistent with an increased need for higher sustained dNTP levels during these conditions.

## Results

### Transposon sequencing reveals synthetic genetic interactions with Lon

We reasoned that pathways regulated by the Lon protease could be identified using a massively parallel mutagenesis strategy such as transposon sequencing, a technique where the number of transposon insertions in a gene of interest reflects the fitness of that gene for a given condition. We generated two biologically independent Tn5 libraries for both NA1000 (wildtype) and *Δlon* strains, plated on nutrient rich media, pooled >200,000 colonies for each library, sequenced transposon insertion sites, compared number of insertions per gene and identified genes where transposon insertions were significantly different (FDR < 0.001) dependent on the presence of Lon (**Figure 1 A, Table S1**).

**Figure 1.**
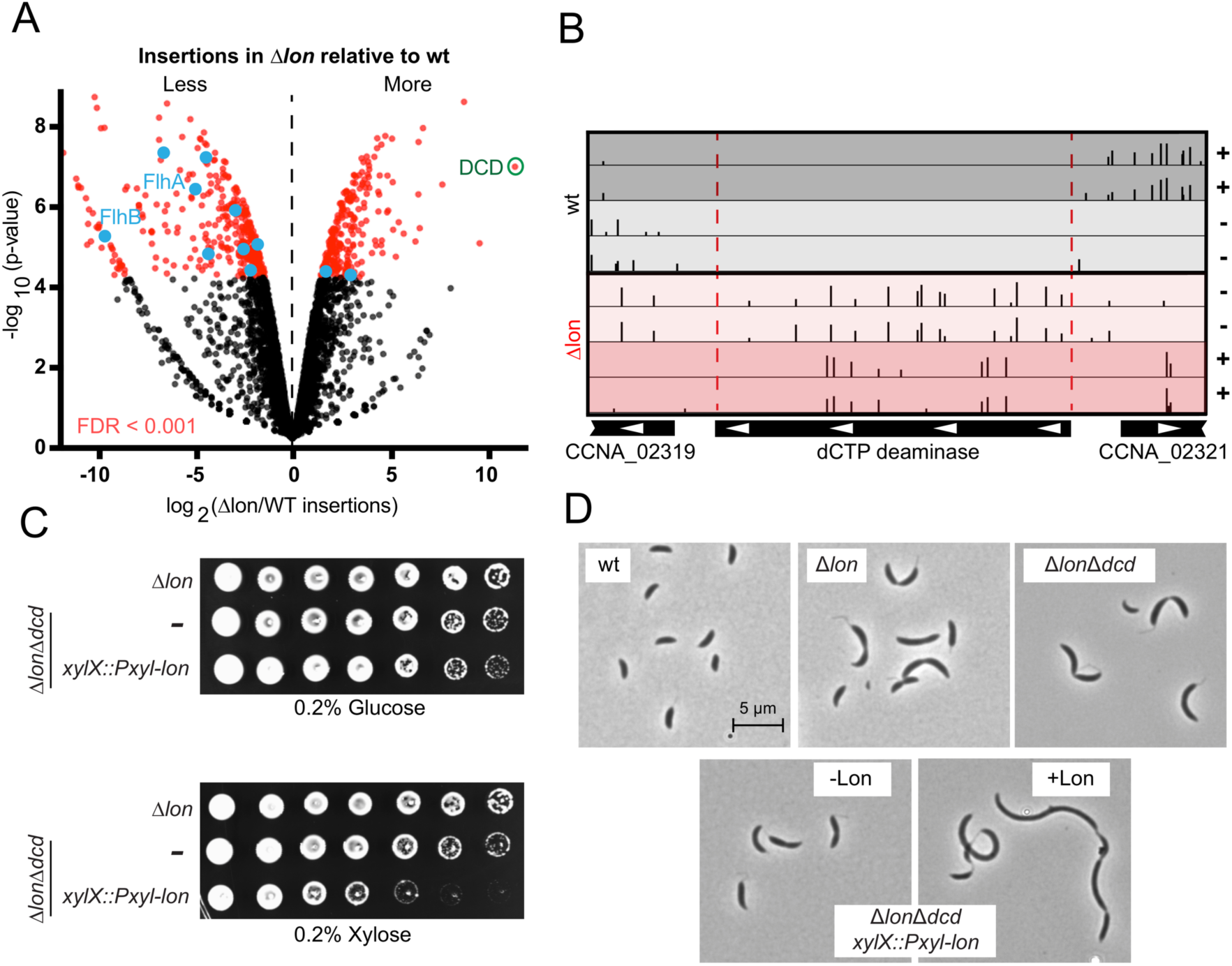
Loss of Lon results in dCTP deaminase dispensability. (A) Volcano plot representation of the Tn-seq analysis of *Δlon* vs wildtype. Negative log_10_ of the p-value is plotted against log_2_ of the fold change in total transposon insertion counts in each gene in wt and *Δlon*. Genes that are either synthetic lethal or conditionally dispensable (FDR <0.001) are highlighted in red. Dark blue represents genes related to flagella assembly and green is CCNA_02320 (dCTP deaminase). (B) Plot of transposon insertion frequency and distribution of dCTP deaminase in wildtype (top) and *Δlon* (bottom) strains. For both wildtype and *Δlon* the positive and negative strand insertions are displayed for both biological replicate transposon libraries indicated by a (+) or (-). The 5’ insertion sites are displayed on a logarithmic scale. (C) Induction of *lon* from the *xylX* promoter results in cell death in the dCTP deaminase deletion (DCD) strain. Cells were grown to exponential phase before being serially diluted 10-fold, and spotted onto media supplemented with 0.2 % glucose (-Lon induction) or 0.2% xylose (+Lon induction). Images of plates were taken on day 3 of growth. (D) Induction of *lon* in from the *xylX* promoter results in filamentation in the dCTP deaminase deletion (DCD) strain. Representative morphologies of indicated strains grown in logarithmic phase. See also Figures S1 and S2 and Table S1.

Kyoto encyclopedia of genes and genomes (KEGG) pathways analysis identified a number of pathways affected, the strongest of which was flagellar synthesis (**Figure 1A, S1**). For example, class II genes related to flagella assembly (Aldridge and Hughes, 2002; Ardissone and Viollier, 2015) had less transposon insertions in general for the Δ*lon* strain versus wildtype (**Figure 1 A, S2**). We confirmed this trend by generating deletions of *flhA* and *flhB* using traditional two-step recombination techniques. We found that strains lacking only *flhA* or *flhB* were easily generated and showed only minor elongation phenotypes, but deletion of these genes in Δ*lon* resulted in severe filamentation and lysis when grown in nutrient rich conditions (**Figure S2**). While finding a link between flagella assembly and Lon was intriguing, we were not surprised about synthetic fitness defects as Δ*lon* strains are already compromised in growth and stress tolerance (Gottesman et al., 1981; Jonas et al., 2013; Wright et al., 1996).

More interestingly, we also identified genes that were more easily disrupted by transposons in cells lacking Lon (**Figure 1 A, S1**). In the strongest case of these epistatic relationships, we identified genes that are annotated as essential in wildtype cells (Christen et al., 2011) but carried many transposon insertions in cells lacking Lon. One such gene was *CCNA_02320* (dCTP deaminase; DCD) which had a well distributed insertion profile across the entirety of its coding sequence only in the Δ*lon* strain (**Figure 1B)**. In support of this observation, a deletion of DCD was easily generated in a Δ*lon* strain, but we could not generate a deletion in the wildtype, consistent with its annotation as an essential gene. Importantly, when we induced Lon in the Δ*dcd*Δ*lon* strain, cells rapidly lost viability and became filamentous (**Figure 1 C,D)** Together, these results show that DCD is “conditionally essential” only when Lon is present, but can be readily deleted in the absence of Lon.

### Loss of Lon elevates deoxynucleotide pools which protects against DCD deletion

By sequencing suppressors which arose upon restoring Lon in a Δ*dcd*Δ*lon* strain, we identified isolates with single mutations in CTP synthetase (CtpS), which converts UTP to CTP (**Figure S3 A,B**). Based on a homology model of *Caulobacter* CtpS these mutants would be predicted to affect substrate binding and likely result in an increase in UTP/CTP ratio (**Figure S3 C)** (Endrizzi et al., 2004). We reasoned that the Δ*lon* strain may tolerate loss of DCD because of changes in nucleotide production and performed metabolomic analysis to investigate this hypothesis.

The metabolic profile of the Δ*lon* strain compared to the parent strain revealed significant differences for 94 metabolites (n=3, FDR <0.1) with many nucleotide related metabolites in this set, including intact molecules and breakdown products (**Figure 2 A and Table S2**).

In general, deoxynucleotides and their breakdown products were elevated in the *Δlon* strain and levels of the equivalent ribonucleotides are reduced, with the exception of dGTP/dGDP (**Figure 2 B**). Interestingly, the metabolite most enriched in the *Δlon* strain is thymine with thymidine and related dTNPs are also elevated (**Figure 2 A,B**).

**Figure 2.**
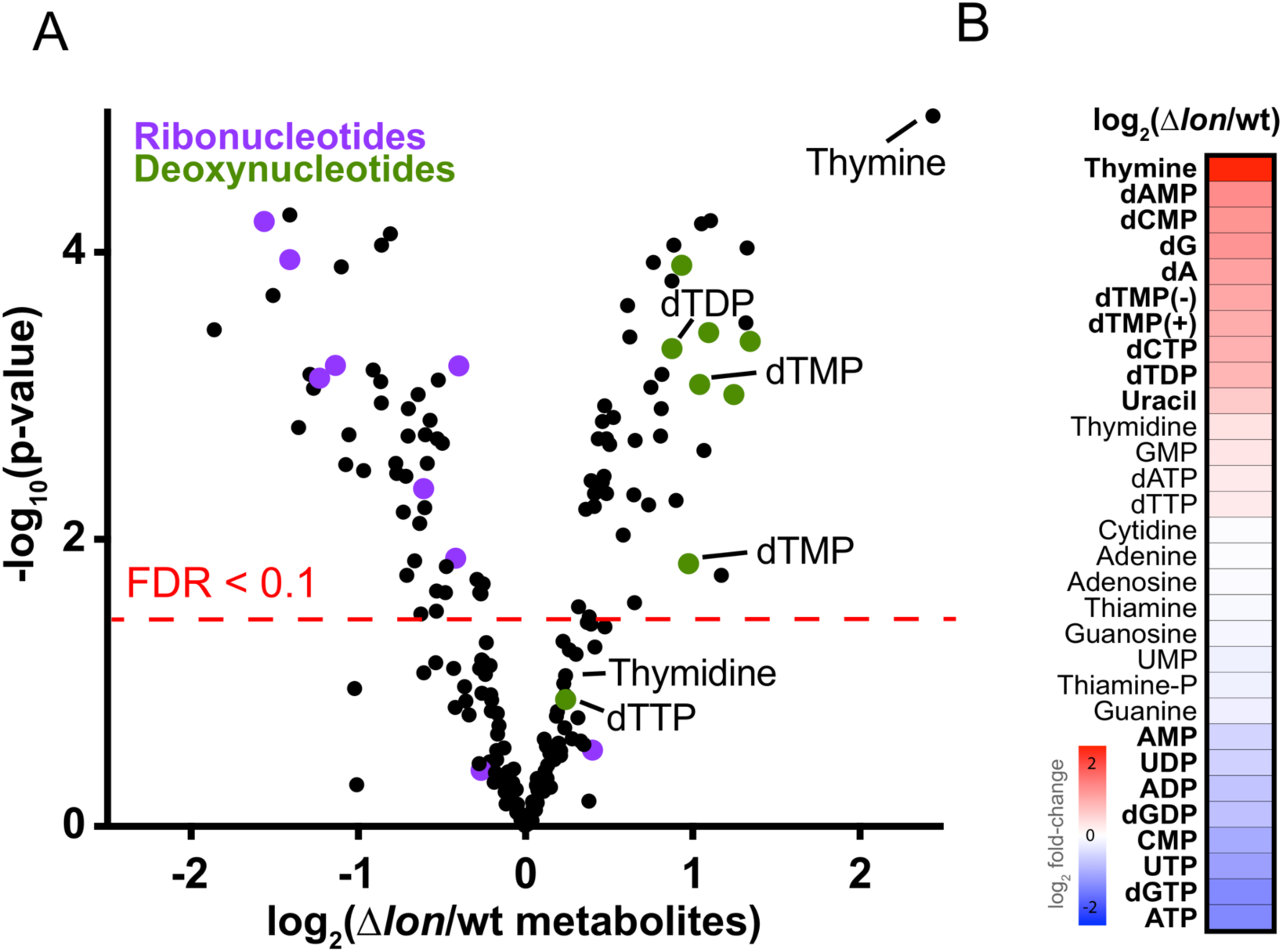
Defects in the *Δlon* strain are associated with metabolic imbalances in nucleotide pools. (A) Volcano plot showing the differences of metabolite levels among the *Δlon* strain compared to wt, as measured by LC-MS. Fold differences were calculated from biological triplicates, run in technical triplicates, and averaged. Cells were grown on 0.22 µm membrane filters placed on top of PYE agar for 4 hours prior to harvesting metabolites. The red dashed line denotes significance (FDR < 0.1). Purple represents ribonucleotides, green deoxynucleotides, and thymidine related metabolites are illustrated. (B) Heatmap showing differences in levels of all nucleotide/precursors identified by LC-MS. Red, White and Blue indicate relative fold difference of increased, neutral, or decreased metabolites in the *Δlon* strain compared to wt. Metabolites with an FDR < 0.1 are bolded. See also Table S2.

DCD catalyzes the deamination of dCTP to dUTP which is dephosphorylated, methylated, and rephosphorylated to generate dTTP (**Figure 3 A**). Therefore, a simple explanation for why loss of DCD is tolerated in the Δ*lon* strain is because dTTP pools are elevated, as supported by our metabolomics data. A corollary to this would be that increasing dTTP in the wildtype strain should protect from loss of DCD. As *Caulobacter* can readily take up exogenous deoxyribonucleotides from the media (Schmidt and Samuelson, 1972; Wagner et al., 2006), we were able to directly test this hypothesis.

**Figure 3.**
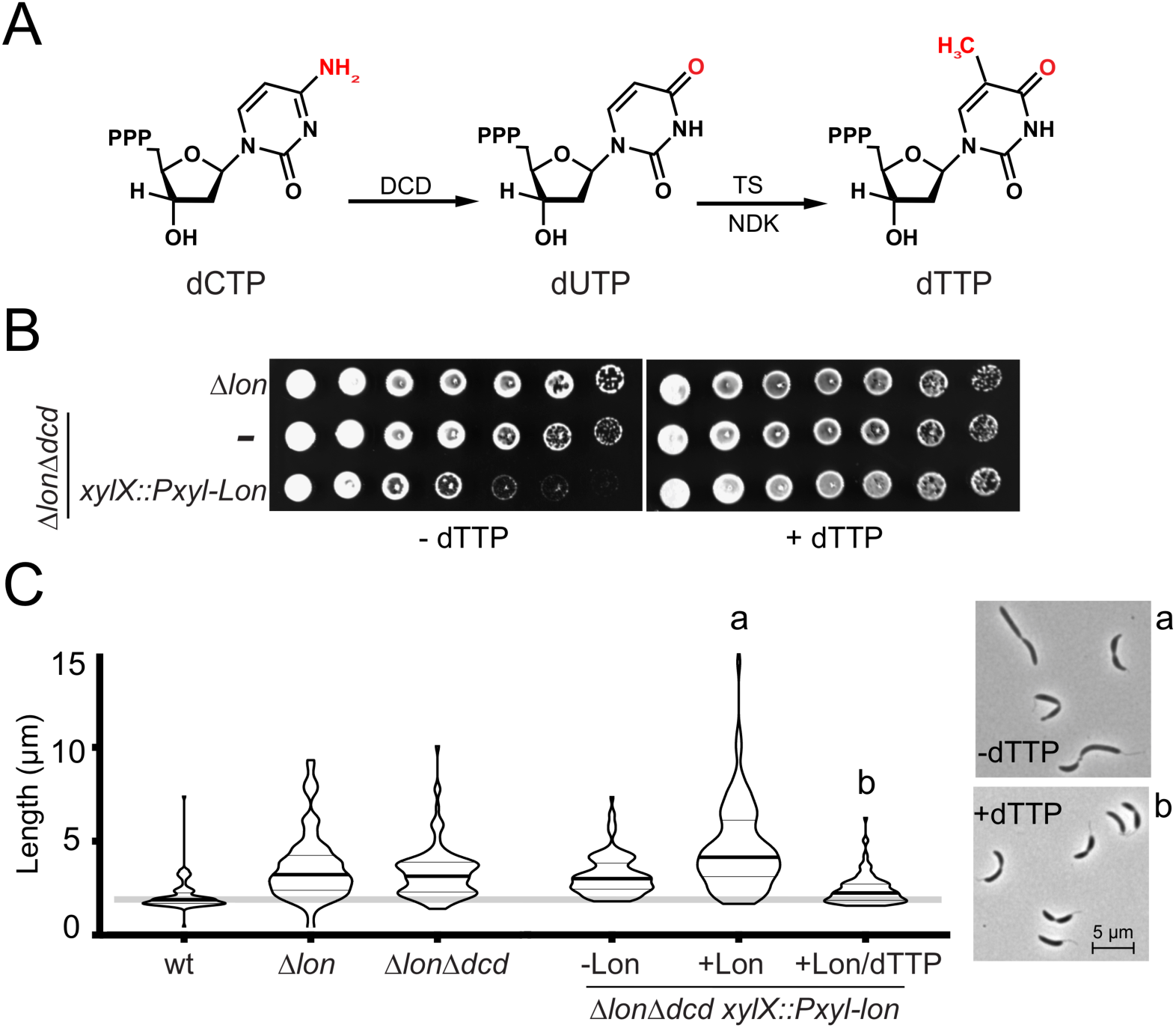
dCTP deaminase is essential for production of dTTP. (A) Nucleotide conversion of dCTP into dUTP by dCTP deaminase (DCD). dUTP can further be converted into dTTP by thymidylate synthetase (TS) and nucleoside-diphosphate kinase (NDK). (B) Cell death induced upon *lon* induction in a dCTP deaminase deletion can be rescued by addition of 100 µM dTTP into solid media. Cells were grown to exponential phase, serially diluted 10-fold, and spotted onto media supplemented with xylose (0.2%; +Lon induction) with or without dTTP addition. (C) Filamentation induced upon *lon* induction in a dCTP deaminase deletion strain can be rescued by addition of dTTP. Cells were outgrown for 7 hours in media containing xylose (0.2%; inducer) or glucose (0.2%; carbon source control) supplemented with or without 100 µM dTTP before being imaged. Quantification of n > 100 with mean and standard deviations. Representative morphologies *of* Δ*lon*Δ*dcd xylX::Pxyl-lon* under inducing conditions (0.2% xylose) with (a) or without (b) supplementation of 100 µM dTTP. See also Figure S3 and S4.

While Δ*dcd*Δ*lon* strains die upon reintroduction of Lon, addition of dTTP to the media restores viability (**Figure 3 B**), as does the addition of dUTP, an intermediate of DCD-dependent dTTP formation. Other dNTPs do not restore viability (**Fig S4).** Addition of exogenous dTTP also prevents the terminal cell filamentation normally seen when Lon is restored to Δ*dcd*Δ*lon* (**Figure 1 C**). We conclude that wildtype cells cannot tolerate a loss of DCD because they cannot maintain appropriate levels of dTTP production in the absence of this pathway.

### *Δlon* have elevated RNR synthesis and activity

Because *Δlon* strains exhibit elevated dTTP levels that allows them to tolerate loss of DCD, we reasoned that other pathways for dNTP production may be upregulated in the absence of Lon. Consistent with this reasoning most deoxynucleotide metabolites are higher in the absence of Lon (**Figure 2 A**). In all cells, ribonucleotide reductase (RNR) generates dNTPs from ribonucleotides and RNA-seq experiments showed increased expression of both RNR subunits (NrdA, NrdB) in the *Δlon* strain, but no significant change in other enzymes related to dTTP production (**Figure 4 A**). Interestingly, we had previously found that although Δ*lon* strains are generally very sensitive to DNA damaging agents (**Figure S5**) (Gottesman et al., 1981; Zeinert et al., 2018), they are 100-fold more resistant to hydroxyurea (**Figure 4 B**). Because hydroxyurea is known to be a direct inhibitor of RNR, our findings that RNR is elevated in Δ*lon* provides a simple explanation for this result. This finding also provides a useful chemical probe to indirectly assess RNR activity using tolerance to hydroxyurea as a proxy.

**Figure 4.**
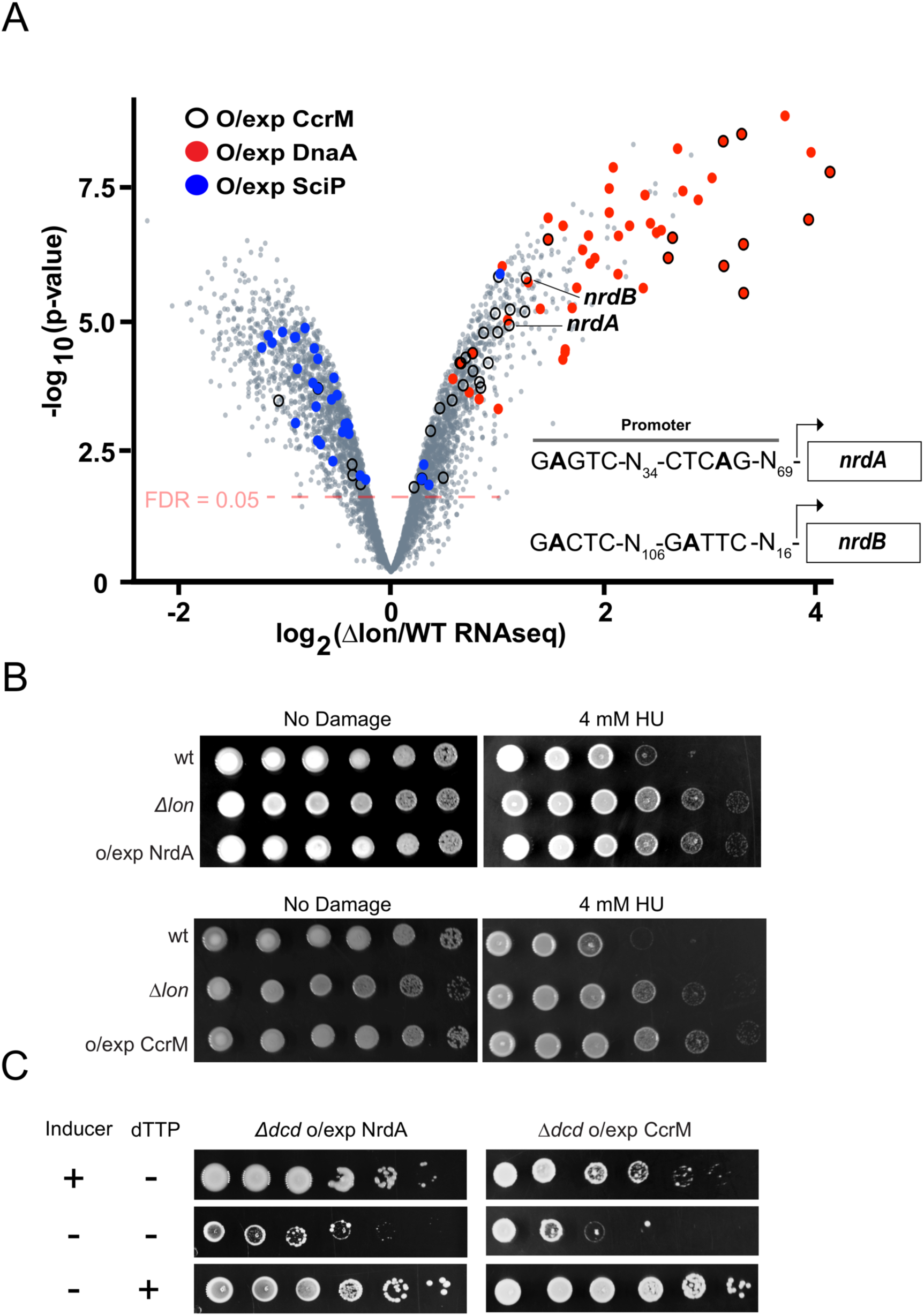
*Δlon* has elevated RNR levels because of CcrM. (A) Volcano plot showing the transcriptional differences of the *Δlon* strain compared to wt, as measured by RNAseq. Negative log_10_ of the p-value is plotted against log_2_ of the fold change mRNA counts in *Δlon* vs wt. Cells were grown to exponential phase before mRNA extraction. The red dashed line denotes significance (FDR < 0.05). Displayed are the top 100 differentially expressed genes as a result of DnaA (red), CcrM (clear black circles), or SciP (blue) overexpression. Illustrated are *nrdA* or *nrdB* promoters and putative CcrM methylation motifs. (B) The *Δlon* mutation and NrdA/CcrM over-expression results in increased resistance to the RNR inhibitor hydroxyurea (HU). Cells were grown to exponential phase before being serially diluted 10-fold, and spotted onto media supplemented with 0.2% xylose (NrdA/CcrM over-expression) with the indicated amount of HU. (C) DCD is dispensable when *nrdA/ccrM* is over-expressed. Cells were grown to exponential phase, serially diluted 10-fold, and spotted onto media supplemented with 0.2% xylose (+ *nrdA* or *ccrM* induction), 0.2% glucose (- *nrdA* or *ccrM* induction), with or without 100 µM dTTP addition. Images of plates were taken on day 4 of growth. See also Figure S5 and Table S3.

Consistent with a protective role for elevated RNR in hydroxyurea resistance, overexpression of NrdA (encoding the alpha subunit of RNR), makes wildtype cells more resistant to hydroxyurea (**Figure 4 B**). If increased RNR levels in the Δ*lon* strain are sufficient to explain DCD dispensability, overexpression of subunits should also support DCD deletion in otherwise wildtype cells. To test this, we deleted DCD in cells overexpressing NrdA and found that these strains are viable, but die when NrdA levels are lowered to normal unless additional dTTP is supplied (**Figure 4 C**). Taken together, these data show that the elevated RNR activity in Δ*lon* is sufficient to provide both protection against hydroxyurea and dispensability of DCD.

### Stabilization of the Lon substrate CcrM transcriptionally upregulates RNR

Next, we explored the mechanism by which Lon controls RNR activity using RNA-seq. Because Lon is a protease, we considered the possibility that Lon degrades a transcriptional activator that controls RNR expression. Two known substrates of Lon, DnaA and CcrM, can be activators of transcription (Gonzalez et al., 2014; Hottes et al., 2005). In *Escherichia coli*, DnaA is known to positively regulate RNR expression (Gon et al., 2006); however, our RNA-seq experiments showed that upregulation of DnaA did not affect RNR expression in *C. crescentus* **(Figure 4A**). CcrM is an adenine methyltransferase that methylates promoter regions in a cell cycle dependent manner (Kozdon et al., 2013; Wright et al., 1996). A global analysis found that the promoter region of NrdA is likely methylated by CcrM and NrdA expression is reduced in the absence of CcrM (Gonzalez et al., 2014). Overexpression of CcrM induced expression of both subunits of RNR (NrdA and NrdB) consistent with the role of CcrM as a transcriptional activator of RNR (**Figure 4A**).

We next tested if elevated CcrM underlies the resistance of Δ*lon* to hydroxyurea. As predicted by our model, overexpression of CcrM in wildtype cells increased hydroxyurea resistance similar to that observed in Δ*lon* (**Figure 4 B**). In addition, CcrM overexpression supports growth of otherwise wildtype cells upon DCD deletion (**Figure 4 C**). Importantly, this strain dies when CcrM levels are lowered to normal levels unless dTTP is supplemented (**Figure 4 C**). Taken together, these data show that a Lon-CcrM-RNR circuit can control dNTP production.

### Misfolded protein burden elevates dNTP pools in a Lon/RNR dependent manner

Lon is well known as a quality control protease responsible for the degradation of bulk misfolded proteins that arise during PQC stress (Goff et al., 1984). Our Lon-CcrM-RNR circuit model suggests that a reduced ability of Lon to degrade CcrM results in elevated dNTP pools via RNR activation. The most extreme case of this activation is seen when Lon is deleted; however, we considered that under PQC stress conditions, Lon degradation of CcrM would be titrated by misfolded protein substrates, leading to a stabilization of CcrM and increased RNR activity.

We first tested whether misfolded proteins could compete for CcrM degradation using a purified system. We performed *in vitro* degradation assays with a well characterized unfolded substrate consisting of a carboxymethylated domain of titin-I27 fused to a hydrophobic region derived from β-galactosidase (CM-titin) (Gur and Sauer, 2008; Jonas et al., 2013) together with purified CcrM and Lon. While Lon alone robustly degrades CcrM on its own with a half-life of ~25 mins, addition of CM-titin results in marked stabilization of CcrM (**Figure 5 A**). We next explored competition *in vivo* by asking if CcrM degradation could be stable in conditions that induced PQC stress. Indeed, growth at an elevated temperature (37°C) resulted in a substantial stabilization of CcrM degradation (**Figure 5 B**). These data suggest that unfolded substrates are capable of competing with natively folded substrates for a limited pool of Lon.

**Figure 5.**
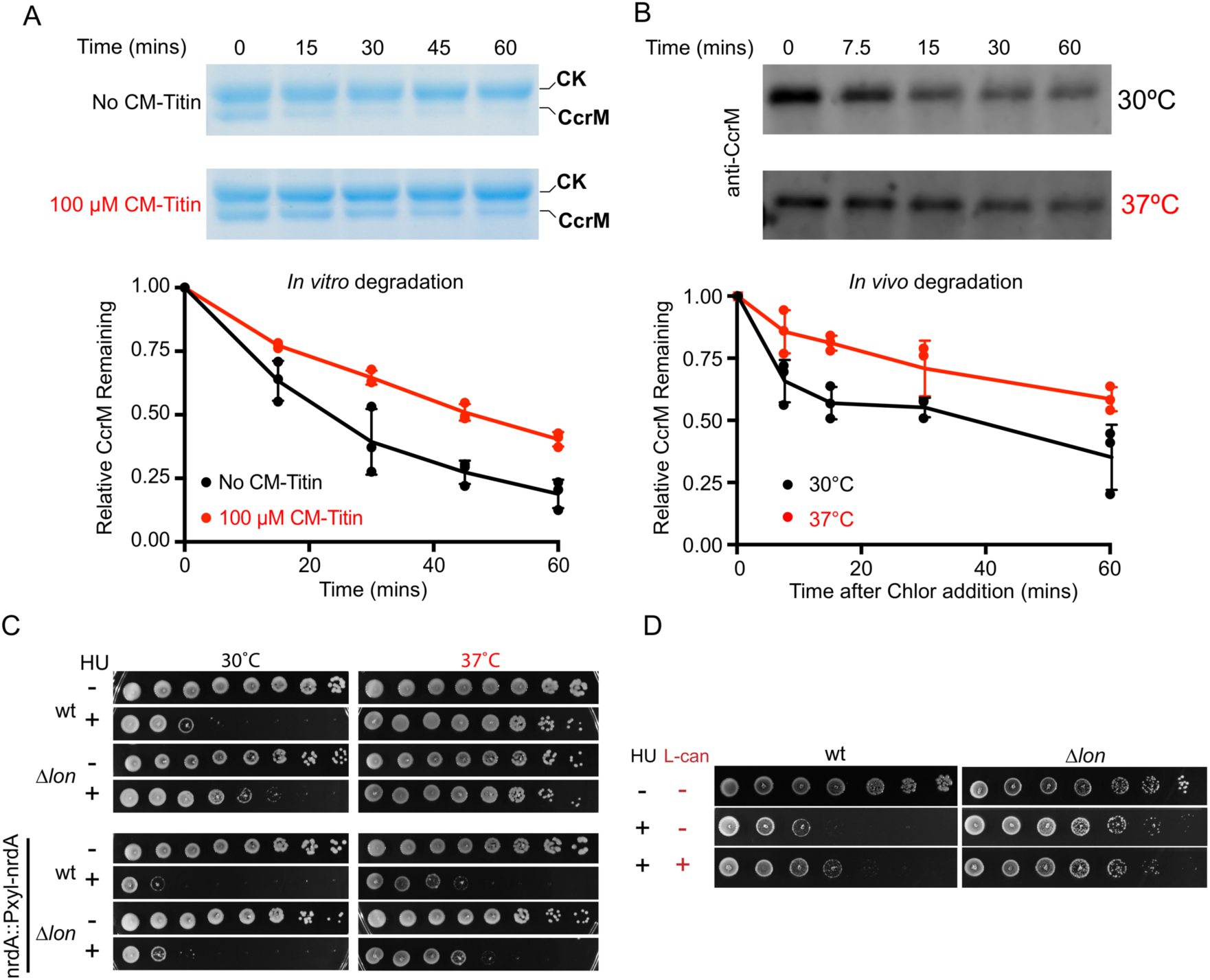
Misfolded protein burden elevates dNTP pools in a Lon/RNR dependent manner. (A) Unfolded substrate (CM-titin) can saturate Lon stabilizing natively folded CcrM *in vitro*. Degradation assays were performed using either CcrM alone or with CM-titin. Reactions consisted of 0.5 µM CcrM, 0.1 µM Lon hexamer, and the indicated concentrations of CM-titin. Quantification of replicate experiments performed in triplicate. Mean and standard deviations plotted with individual data points shown. (B) Growth at elevated temperatures increases CcrM stability *in vivo*. Cells were grown to exponential phase at 30 or 37°C. Protein synthesis was inhibited by addition of chloramphenicol. Samples were withdrawn at the indicated time points and immediately quenched in SDS lysis buffer. Lysate from equal number of cells was used for Western blot analysis and probed with anti-CcrM antibody. A cropped representative image and quantification of biological triplicates is shown. Error bars represent standard deviation. (C) Thermal stress increases resistance to hydroxyurea in a Lon dependent manner. Strains were grown to exponential phase before being serially diluted 10-fold and spotted onto media supplemented with 0.2% xylose for *nrdA* induction, with or without 5 mM hydroxyurea (HU). Plates were then grown for 2 days at 37°C or 3 days at 30°C before imaging. (D) Addition of proteotoxic stress increases resistance to hydroxyurea in a Lon dependent manner. Strains were grown to exponential phase before being serially diluted 10-folded, and spotted onto PYE media supplemented with 5 mM hydroxyurea in combination with 50 µg/mL L-canavanine (L-can).

Because stabilization of CcrM leads to hydroxyurea resistance due to increased RNR activity, we next asked whether increased PQC stress could also lead to similar effects due to titration of Lon activity by misfolded proteins. Interestingly, when wt cells were grown at 37°C compared to 30°C there was a marked increase in hydroxyurea resistance (**Figure 5 C**). The Δ*lon* strain was more resistant to hydroxyurea at 30°C and increased temperature didn’t elicit as strong of a protective effect as what was observed in the wt strain. Similar Lon-specific protection against hydroxyurea was seen when L-canavanine was used to induce PQC stress (**Figure 5 D**). To specifically test if this effect was due to the Lon-CcrM-RNR circuit described above, we generated a strain where the only copy of NrdA was expressed under the control of the xylose inducible promoter (*nrdA::Pxyl-nrdA*), thereby decoupling the Lon-CcrM-RNR circuit as the xylose promoter is not under CcrM control (Meisenzahl et al., 1997; Thanbichler et al., 2007). These strains were fully viable in the absence of hydroxyurea at both 30°C and 37°C, but there was no longer any Lon-specific protection at elevated temperature (**Figure 5 C**). Taken together, these data show that PQC stress *in vivo* is sufficient to induce a Lon-dependent increase in RNR activity through CcrM stabilization.

### Competition sustains growth during increased misfolded protein burden

Finally, we explored whether this PQC responsive Lon-CcrM-RNR circuit was important for growth without additional stresses such as hydroxyurea. We reasoned that if our Lon titration model was important for responding to PQC stress, then decoupling the CcrM-dependent control of NrdA by using the *nrdA::Pxyl-rdA* strain would result in reduced fitness during stress. We used competition assays to determine fitness (**Figure 6 A**), monitoring growth of fluorescently labeled wildtype strains in competition with unlabeled *nrdA::Pxyl-nrdA* strain. We found that at 30°C, both wildtype and *nrdA::Pxyl-nrdA* strains competed equally well. However, at 37°C there was a defect in competitive fitness of the *nrdA::Pxyl-nrdA* strain (**Figure 6 B**). This defect became more pronounced when levels of *nrdA* were reduced showing even stronger fitness defects upon comparing 30°C vs 37°C. We conclude that stabilization of CcrM upon titration of Lon by PQC stress (misfolded proteins) modulates dNTP pools which is important for adaptation to that stress.

**Figure 6:**
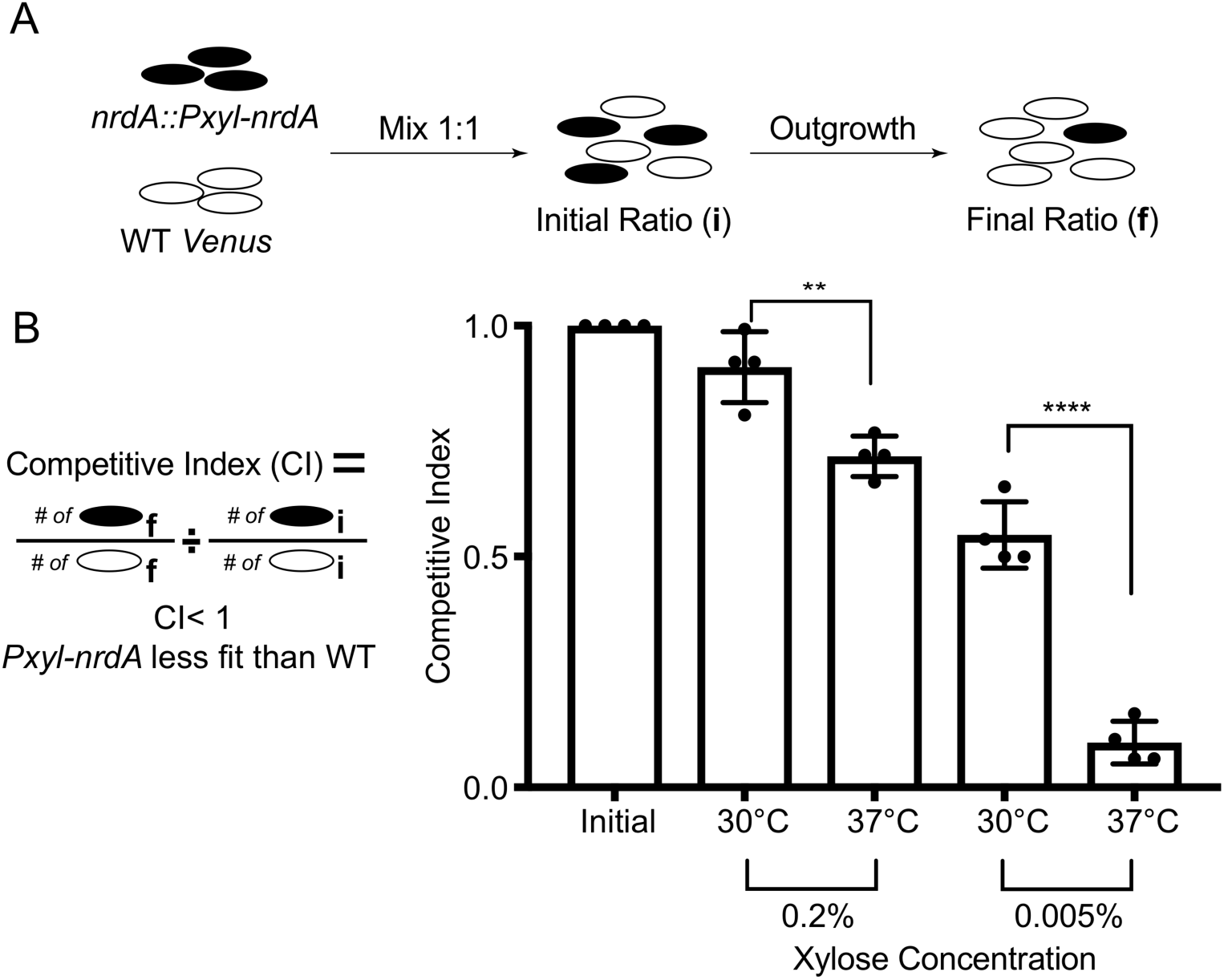
CcrM-dependent control of NrdA is protective for growth at elevated temperatures. (A) Schematic of assay to assess fitness. (B) A strain in which *nrdA* is transcriptionally controlled by the *xylX* promoter has reduced fitness during growth at elevated temperatures. Competition assay with *xylX::Plac-venus* (constitutive venus expression) and *nrdA::Pxyl-nrdA*. Cells were mixed at one to one ratio, diluted 1:500, and outgrown in the respective conditions. Quantification of n > 100 cells. Mean and standard deviation included for bar graphs as well as performed a student’s t-test using two tails. A * indicates significance of p< 0.05.

## Discussion

Lon was one of the first energy-dependent proteases identified (Charette et al., 1981) and is principally thought of as a protein quality control (PQC) protease, degrading misfolded or damaged proteins (Goff et al., 1984; Gottesman and Zipser, 1978). In *E. coli*, Lon is known to degrade cell division inhibitors and capsule regulators (Mizusawa and Gottesman, 1983; Torres-Cabassa and Gottesman, 1987) that reflect the role of Lon in damage responses and bacteria development. Indeed, loss of Lon in many species results in an inability to response to stress conditions (Ching et al., 2019; Jonas et al., 2013; Ngo and Davies, 2009; Robertson et al., 2000), particularly those that induce misfolded proteins (Goff and Goldberg, 1987; Goff et al., 1984; Jonas et al., 2013). However, despite the importance of this protease, specific pathways that are influenced by Lon during both normal and stress conditions are poorly understood in most bacteria, such as *Caulobacter crescentus*.

In this study, we found that levels of dNTPs are governed by Lon dependent degradation of the DNA adenine methyltransferase CcrM and that this regulation is especially important during conditions which increase the misfolded protein burden (**Figure 7**). We used transposon sequencing to broadly identify genetic interactions with the loss of Lon. One of the strongest of these synthetic interactions was that the normally essential dCTP deaminase was now dispensable in Δ*lon*. Metabolomic profiling showed that overall levels of dNTPs are elevated in Δ*lon*, including dTTP, the terminal product of dCTP deamination. Consistent with this, dCTP deaminase could be deleted in otherwise wildtype cells by adding dTTP exogenously. We linked the higher dNTP production in Δ*lon* to an increase in ribonucleotide reductase expression driven by stabilization of the transcription factor CcrM, a known Lon substrate. The increased ribonucleotide reductase in Δ*lon* is consistent with the exceptional resistance of these strains to hydroxyurea, a result that was originally confusing to us given the extreme sensitivity of Δ*lon* to many other DNA damaging stresses (**Figure S5**) (Zeinert et al., 2018). We found that this increased resistance to hydroxyurea was also seen in wildtype strains during protein misfolding conditions, suggesting that competition for a limited pool of protease could result in increased stability of Lon substrates during stress. Indeed, Lon degradation of CcrM can be titrated away by misfolded proteins both *in vitro* and *in vivo*. We propose that this competition mechanism allows cells to flexibly respond to a variety of many protein damaging conditions by activating CcrM dependent pathways, such as increased dNTP pools, sensed through the increased burden on Lon.

**Figure 7.**
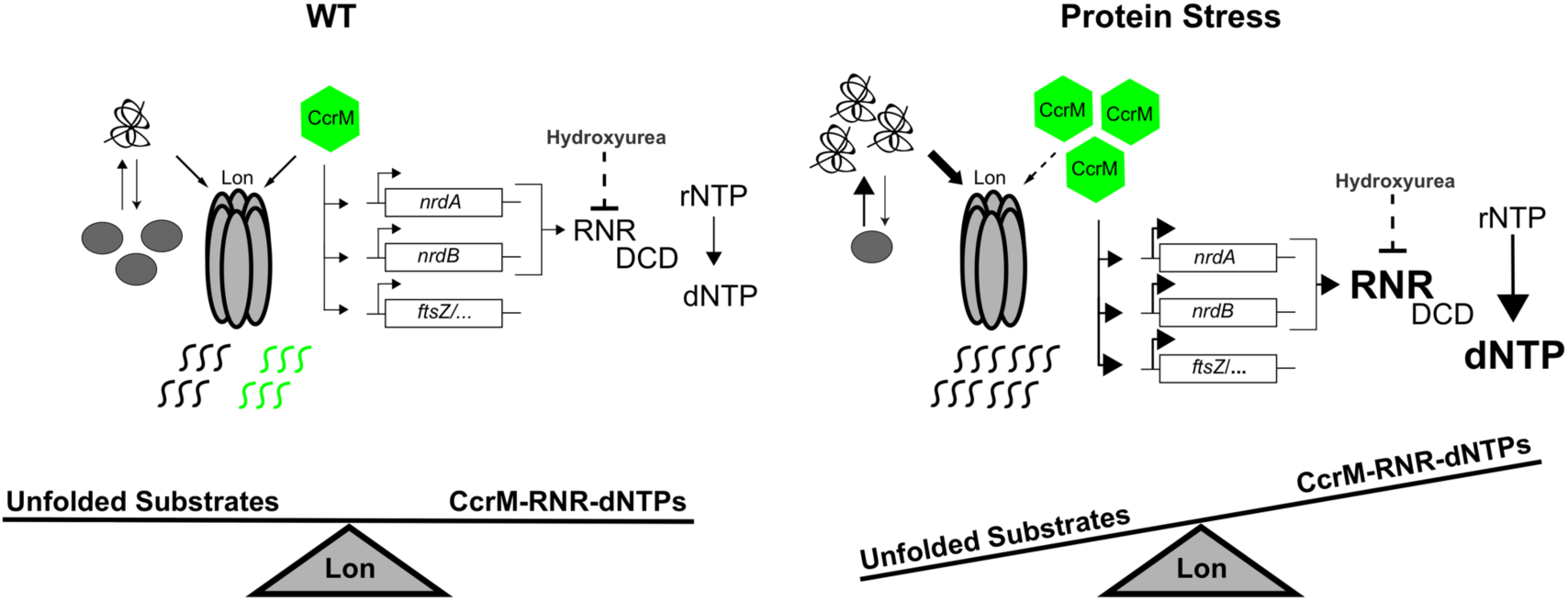
Model for Lon-CcrM-RNR circuit. In wildtype cells Lon restricts dNTP pools through degradation of the adenine methyltransferase CcrM keeping transcription of *nrdA*/*nrdB* relatively low which puts selective pressure on dCTP deaminase (DCD) dependent conversion of dCTP into dTTP. In strains lacking Lon or taxed quality control, CcrM levels are elevated resulting in constitutive promoter methylation of *nrdA*/*nrdB* and consequently increased transcription. Increased levels of RNR levels removes selective pressure from DCD allowing for its dispensability. Strains possessing elevated CcrM have increased levels of RNR making them more resistant to the drug hydroxyurea. Lon integrates its functions in quality control with those of native substrate degradation. This presents competition between unfolded and natively folded substrates allowing for growth under conditions that result in protein misfolding.

Why would an increased dNTP pool be beneficial during stress conditions? We speculate that this would be most useful when increased protein synthesis results in increased misfolded protein burden, such as we see with rapid growth at elevated temperature. It is also possible that this circuit plays an important role when Lon substrates other than misfolded proteins are upregulated, such as during DNA damage or oxidative stress. For DNA damage it is interesting to consider that sufficient dNTP levels are important for fueling the cycles of repair/replication needed to fix most types of DNA defects.

Titrating activity of limited PQC proteins is an elegant strategy for cells to respond to the products of stress conditions rather than the specific initiating signal. In bacteria, the heat stress sigma factor sigma32 is bound to the DnaK chaperone and is thought to be released when DnaK becomes saturated by increased misfolded proteins during heat stress (Arsène et al., 2000; Blaszczak et al., 1999). Similarly, in eukaryotes, titration of Hsp70 by misfolded proteins results in release of active Hsf1, the major proteostasis transcription factor (Masser et al., 2019). Titration of bacterial proteases has also been shown to control some responses, for example, in *E. coli* upregulating of RpoS during translational stress is attributed in part to accumulation of ClpXP substrates resulting from oxidative damage (Fredriksson et al., 2007). In synthetic biology this titration of proteases has been used to generate delays and coupling in gene circuits through a process called protease queueing (Prindle et al., 2014). Finally, in *E. coli* overexpression of the Lon substrate SulA can indirectly stabilize another substrate RcsA, suggesting that Lon titration is not limited to PQC stresses (Dervyn et al., 1990).

The work we have presented here is a natural biological circuit that integrates a variety of protein stresses through titration of the Lon protease to ensure cell fitness during these conditions. Given the conservation of Lon in bacteria and its general ability to recognize misfolded proteins, we suspect that that similar connections will exist between PQC and growth promoting cell processes in other bacteria. Because Lon substrates are also upregulated during other damaging conditions, such as DNA damage (Mizusawa and Gottesman, 1983), we predict that a Lon titration circuit likely plays roles in stress responses beyond PQC.

## Methods

### Contact for Reagent and Resource Sharing

Question about or requests for methods, strains, and reagents should be directed to and will be fulfilled by the Lead Contact, Peter Chien.

### Experimental Model and Subject Details

#### Growth conditions and chemical treatments

*Caulobacter crescentus* NA1000 (wt) strains were cultured in peptone (2 g/L) yeast (1g/L) extract medium (PYE) supplemented with 1 mM MgSO_4_, 0.5 mM CaCl_2_, and 1.5% agar at 30 °C unless otherwise indicated. For induction purposes, the Pxyl promoter was induced with xylose (0.2%) and repressed with glucose (0.2%). To generate strains, antibiotics were added at the following concentrations (plates): kanamycin (25 μg/ml), gentamycin (5 μg/ml), spectinomycin (100 μg/ml), and oxytetracycline (2 μg/ml). After initial selection steps for strain construction, antibiotics were excluded from all liquid and solid media assays. *E. coli* strains were grown in LB (10 g/L NaCl, 10 g/L tryptone, 5 g/L yeast extract) and supplemented with antibiotics at the following concentrations (liquid/plates): oxytetracycline (15/15 μg/ml), spectinomycin (50/50 μg/ml), kanamycin (50/50 μg/ml), gentamycin (20/20 μg/ml). For recombinant protein expression, 0.4 M IPTG was used to induce gene expression. Optical density was measured at 600 nm.

#### Strain construction

All *Caulobacter* strains were derivatives of CPC176, an isolate of strain CB15N/NA1000. Deletion strains were constructed using the pNPTS138 plasmids (Skerker et al., 2005) which uses a two-step recombination procedure with counter selection by sucrose/*sacB*. This procedure was followed to produce strains: CPC592, CPC593, CPC594, CPC595 and CPC619. Following transformation with pNPTS_FlhA_KO, pNPTS_FlhB_KO, or pNPTS_DCD_KO into CPC176 or CPC456, initial primary selection was on PYE with kanamycin. Primary colonies were grown overnight without selection to allow for secondary recombination. Overnight cultures were plated onto PYE agar plates supplemented with 3% w/v sucrose and gentamycin. To validate gene deletion, resulting strains were tested for antibiotic sensitivity to kanamycin and gentamycin.

CPC774 and CPC813, were generated by electroporating pNPTS_DCD_KO into CPC739 and CPC812. Following transformation, initial primary selection was on PYE supplemented with kanamycin and xylose. Primary colonies were grown overnight without selection to allow for secondary recombination in the presence of xylose. Overnight colonies cultures were plated onto PYE agar plates supplemented with 3% w/v sucrose, xylose and gentamycin.

For overexpression or depletion, strains were constructed using a one-step recombination procedure at the xylose locus, using pXGFPN-5 or at the endogenous locus, using pXMCS-5 (Thanbichler et al., 2007).

CPC624 was constructed by electroporating plasmid pXGFPN-5_CcLon into CPC619 and selecting onto oxytetracycline plates. CPC624 was validated by anti-Lon western for clones that expressed Lon with xylose and had low levels of Lon with no induction.

CPC739 and CPC812 were generated by electroporating pXGFPN-5_CcrM or pXGFPN-5_NrdA into CPC176 and selecting on oxytetracycline plates. CPC739 was validated by anti-CcrM western for clones that expressed elevated levels of CcrM with xylose and had wt levels of CcrM with no induction. CPC812 was validated by PCR with primers OPC21 and OPC14 and further validated phenotypically using hydroxyurea resistance as an indirect measure.

The NrdA depletion strains, CPC882 and CPC883, were generated by electroporating pXMCS-5_NrdA into CPC456 and selecting on oxytetracycline plates supplemented with xylose. Followed by fresh CR30 phage transduction into recipient strains CPC176 and CPC456 and validated by viability with or without inducer, and sensitivity to hydroxyurea.

#### Plasmid Construction

##### Integration plasmids

All deletion plasmids are comprised of 1000 bp upstream and downstream of the coding regions flanking a gentamycin resistance cassette.

pNPTS_DCD_KO was constructed by amplifying the upstream/downstream region of *DCD* with primers 5_F_UTR_2320 and 5_R_UTR_2320 and 3_F_UTR_2320 and 3_R_UTR_2320. The gentamycin resistance cassette was amplified using the primers F_gentamycin and R_gentamycin. The vector pNPTS138 was digested with HinIII and EcoRI. Isothermal assembly was performed with equal molar concentrations of 5’, 3’, gentamycin resistance, and vector.

pNPTS_FlhA_KO was constructed by amplifying the upstream/downstream region of *DCD* with primers 5_F_UTR_FlhA and 5_R_UTR_FlhA and 3_F_UTR_FlhA and 3_R_UTR_FlhA. The gentamycin resistance cassette was amplified using the primers F_gentamycin and R_gentamycin. The vector pNPTS138 was digested with HinIII and EcoRI. Isothermal assembly was performed with equal molar concentrations of 5’, 3’, gentamycin resistance, and vector.

pNPTS_FlhB_KO was constructed by amplifying the upstream/downstream region of *DCD* with primers 5_F_UTR_FlhB and 5_R_UTR_FlhB and 3_F_UTR_FlhB and 3_R_UTR_FlhB. The gentamycin resistance cassette was amplified using the primers F_gentamycin and R_gentamycin. The vector pNPTS138 was digested with HinIII and EcoRI. Isothermal assembly was performed with equal molar concentrations of 5’, 3’, gentamycin resistance, and vector.

pXGFPN-5_CcLon was constructed by amplifying the coding sequence of Lon with F_Lon_pXGFPN_5 and R_Lon_pXGFPN_5. The vector pXGFPN-5 was digested with NdeI and EcoRI. Isothermal assembly was performed with equal molar concentrations of insert and vector.

pXGFPN-5_NrdA was constructed by amplifying the coding sequence of NrdA with F_NrdA_pXGFPN_5 and R_NrdA_pXGFPN_5. The vector pXGFPN-5 was digested with NdeI and EcoRI. Isothermal assembly was performed with equal molar concentrations of insert and vector.

pXGFPN-5_CcrM was constructed by amplifying the coding sequence of CcrM with F_CcrM_pXGFPN_5 and R_CcrM_pXGFPN_5. The vector pXGFPN-5 was digested with NdeI and EcoRI. Isothermal assembly was performed with equal molar concentrations of insert and vector.

pXMCS-5_nrdA was constructed by amplifying a fragment of NrdA with F_nrdA_XMCS_5 and R_nrdA_truncation_XMCS_5. The vector pXMCS-5 was digested with NdeI and EcoRI. Isothermal assembly was performed with equal molar concentrations of insert and vector.

## Method Details

### Plating viability and drug sensitivity

All *Caulobacter* strains were grown overnight in PYE media. After overnight growth, cells were back diluted to OD_600_ 0.1 and outgrown to mid-exponential phase before being normalized to OD_600_ 0.1 and 10-fold serially diluted on to media lacking carbon addition or containing xylose (O.2%) for induction or glucose (0.2%) for repression. For the dCTP deaminase deletion strains (CPC619 and CPC624) dTTP was supplemented into the media at 10 μM to ensure genomic stability and decrease the chance of spontaneous suppressor mutations. For wt and Δ*lon nrdA::Pxyl-nrdA* strains (CPC882 and CPC883) xylose (0.2%) was added in hydroxyurea damage assays.

For experiments using hydroxyurea (Sigma), mitomycin C (Sigma), methyl methanesulfonate (Sigma) or L-canavanine (Sigma), damaging agents were freshly prepared at a stock concentration of 1M hydroxyurea, 0.4 µg/ml mitomycin C, and 100 mg/ml L-canavanine and filter sterilized. Media was allowed to cool prior to adding final concentrations of 3, 4, and 5 mM hydroxyurea with/without 50 µg/ml L-canavanine, 0.0001% methyl methanesulfonate, and 1 µg/ml Mitomycin C. Agar plates were allowed to air dry 2 hours prior to serially dilution plating. All Plates were incubated at 30°C unless otherwise indicated for 2-3 days and imaged with a gel doc.

For experiments using deoxynucleotide additions: dTTP (Sigma), dUTP (Sigma), and dCTP (NEB). All stocks were freshly prepared at concentrations of 100 mM and filter sterilized. Media was cooled prior to adding final concentrations of 10 or 100 µM dNTP. Agar plates were 2 hours prior to serially dilution plating. Plates were incubated at 30°C for 3 days and imaged with a gel doc.

### *In vivo* growth competition assay

Overnight cultures of a strain constitutively expressing the fluorescent reporter Venus (CPC798) was mixed with *nrdA::Pxy-:nrdA* (CPC882) at a 1:1 ratio. Mixed strains were then diluted 1:500 into fresh media and allowed to outgrow at both 30 and 37°C overnight. The initial 1:1 mixture of cells was verified by phase contrast and fluorescent microscopy. Quantification of >200 cells was performed for each condition and biological replicate. All final ratios were normalized to their starting ratios prior to dilution and outgrowth. Data were plotted using Graphpad Prism version 8.3.0. Statistical analysis of 4n biological replicates consisted of a two-tailed Student t-test. A * denotes p-value < 0.05, ** <0.01, **** <0.0001.

### Microscopy

Microscopy images are phase contrast images of exponentially growing cells (Zeiss AXIO ScopeA1) mounted on 1% PYE agar pads using a 100X objective. Cells lengths were quantified using MicrobeJ (Ducret et al., 2016) for ImageJ (Schneider et al., 2012). Prism was used for graphical representation of cell length measurements. Representative images of the same scale were cropped to illustrate morphological defects. Phase contrast and fluorescent images were taken, over-lay, and manually counted for competition assays.

### Transposon Mutagenesis and Deep Sequencing

For Figure 1, two Tn-seq libraries were generated for both wt and Δ*lon*. 2L PYE cultures were pelleted at OD_600_ 0.4-0.8, washed with 10% glycerol, and electroporated with the Ez-Tn5 <Kan-2> transposome (Lucigen). Cells were recovered for 90 mins at 30 °C with shaking, and plated onto PYE + kan plates. WT libraries were grown for 5 days and 7 days for Δ*lon* libraries. Colonies were scraped from the surfaces of plates, combined, resuspended to form a homogenous solution of PYE + 20% glycerol as a cryoprotectant.

Genomic DNA was extracted from an aliquot of each library using the MasterPure Complete DNA and RNA purification kit according to the manufacturer’s protocol (Epicentre). Libraries were then prepared for Illumina Next-Generation Sequencing via PCR. Final indexed libraries were pooled and sequenced at the University of Massachusetts Amherst Genomics Core Facility on a NextSeq 500 (Illumina).

For Tn-seq, single end 75 base reads were first demultiplexed by index, each library was concatenated and the molecular modifier added in the second PCR was clipped using Je (Girardot et al., 2016). Clipped reads were mapped to the *Caulobacter crescentus* NA1000 genome (NCBI Reference Sequence: NC_011916.1) using BWA (Li and Durbin, 2010), sorted with Samtools (Li et al., 2009). Duplicate transposon reads introduced by PCR were removed using Je (Girardot et al., 2016) and indexed with Samtools (Li et al., 2009). For ease of data visualization, the 5’ insertion sites of each transposon were converted to a wiggle file (.wig) and visualized using Integrative Genomics Viewer (IGV) (Robinson et al., 2011). Specific hits for each library were determined by calculating per position numbers of 5’ sites for unique inserts across the entire genome using *bedtools genomecov* (Quinlan and Hall, 2010), then counting the number of unique 5’ insertions within the middle 80% of each gene using custom bedfiles with truncated ORFs and *bedtools map*. For statistical comparison, a quasi-likelihood F-test (glmQLFit) from the edgeR package in the Bioconductor suite (Robinson et al., 2009) was used to determine the false discovery rate adjusted p-values. Specific scripts are available upon request.

For RNA-seq, exponential phase cells were harvested, RNA was extracted and rRNA was depleted using Ribo-zero (Illumina). Sequencing libraries were generated using NEB Next RNA Library Prep kits and sequenced on a NextSeq 500 with single end 75 base reads. Reads were mapped to the *Caulobacter* NA1000 genome using BWA, sorted with Samtools and the resulting BAM files were used as input with *bedtools map* and a bedfile consisting of the all gene features to count the number of reads per gene. Plots were produced using Rstudio and ggplot (R Core Team, 2019; Wickham, 2016)

### Metabolite Extraction

Metabolite extraction was performed as described previously (Irnov et al., 2017). Briefly, *C. crescentus* strains were grown at 30°C in PYE until reaching OD_600_ 0.4. Approximately 30 mL of cells were transferred onto a sterile 0.22 μm polyethersulfone (PES) membrane filter (Millipore) by vacuum filtration. The filters were then placed on the surface of PYE agar plates and cells were allowed to grow for 4 hours at 30°C. Samples were quenched by placing the filters into acetonitrile/methanol/H_2_O (40:40:20, kept at -20°C). Cells were washed off the membrane using the quenching buffer. The cells were then transferred to -80°C prior to analysis on LC-MS.

### Liquid chromatography-mass spectrometry (LC-MS) analysis

#### LC-MS/MS system

Mass spectrometric analyses were performed on Sciex “QTRAP 6500+” mass spectrometer equipped with an ESI ion spray source. The ESI source was used in both positive and negative ion modes. The ion spray needle voltages used for MRM positive and negative polarity modes were set at 4800 V and -4000 V, respectively. The mass spectrometer was coupled to Shimadzu HPLC (Nexera X2 LC-30AD). The system is under control by Analyst 1.6.3 software.

#### Chromatography Conditions

Chromatography was performed under HILIC conditions using a SeQuant® ZIC®-pHILIC 5 µm polymeric 150 × 2.1 mm PEEK coated HPLC column, MilliporeSigma, USA. The column temperature, sample injection volume, the flow rate was set to 45 °C, 5 µL, and 0.15 mL/min respectively. The HPLC conditions were as follows: Solvent A: 20 mM ammonium carbonate including 0.1% Ammonium hydroxide. Solvent B: Acetonitrile. Gradient condition was 0 min: 80% B, 20 min: 20% B, 20.5 min 80% B, 34 min: 80% B. Total run time: 34 mins. Data was processed by SCIEX MultiQuant 3.0.3 software with relative quantification based on the peak area of each metabolite.

### Protein Purification and Modification

CcrM was affinity purified by addition of His6SUMO to the N-terminus (Wang et al., 2007). His_6_SumoCcrM was expressed in BL21(DE3) pLysS *E. coli*. The cells were grown until OD_600_ (0.8), induced with 0.4 mM isopropyl 1-thio-β-D-galactopyranoside (IPTG) for 12 hours at room temperature without shaking followed by centrifuged at 5,000 RPM for 10 mins. The pellets were resuspended in lysis buffer (50 mM Tris (pH 8.0), 300 mM NaCl, 20 mM imidazole, 10% glycerol, and 1 mM DTT. The cell suspension containing 1 mM PMSF was lysed by microfluidizer (Microfluidics, Newton, MA). The lysate was cleared at 14,5000 RPM for 30 minutes at 4°C. Clarified lysate was loaded onto a pre-equilibrated Ni-NTA resin, washed three times with 3 column volumes lysis buffer and finally eluted. The eluted protein was buffer exchanged into lysis buffer without imidazole and the His_6_SUMO tag was cleaved by Ulp1-his protease for 1 hour at room temperature. The tags were then further removed by subtractive Ni-NTA affinity chromatography. Native Lon protease was purified as previously described (Gur and Sauer, 2008; Liu et al., 2019). The ‘titin’ construct was purified by Ni-NTA as previously described (Gur and Sauer, 2008; Liu et al., 2019). CM-titin was generated by carboxy methylation of two cysteine residues in the titin-I27 with iodoacetamide under guanidine hydrochloride denaturation. Modified protein was buffer exchanged into TK buffer (25 mM Tris pH 8.0, 100 mM KCl, 10 mM MgCl_2_, and 1 mM DTT) and stored at 4°C for stability reasons.

### In vitro degradation assay

Degradation for all reactions were performed at 30°C with the following protein concentrations unless elsewhere indicated: 0.1 μM Lon_6_, 0.5 μM CcrM, CM-titin (100 and 50 μM), with 4 mM ATP, 15 mM creatine phosphate (Sigma) and 75 μg/mL creatine kinase (Roche) as ATP regeneration components. The reactions were initiated by adding the ATP regeneration mix to the protease-substrate solution in TK buffer (25 mM Tris pH 8.0, 100 mM KCl, 10 mM MgCl_2_, and 1 mM DTT). Twenty μL aliquots were taken at each indicated time point and quenched in SDS loading dye (2% SDS, 6% Glycerol, 50 mM Tris pH 6.8 and 2 mM DTT), and examined by 12% SDS-PAGE gels. Densitometry was performed with imageJ (Schneider et al., 2012), band intensities were normalized using creatine kinase as an internal loading control and degradation rates were plotted using Prism.

### *In vivo* protein stability assays

Wildtype strains were grown overnight in PYE media at either 30°C or 37°C. After overnight growth, cells were back diluted to OD_600_ 0.1 and outgrown to mid-exponential phase at their respective temperatures. Protein synthesis was blocked by addition of 30 µg/ml chloramphenicol. Cell samples were removed at the indicated time points and immediately lysed in SDS lysis buffer and stored at -20°C prior to western blot analysis.

### Western Blot Analysis

Cell cultures withdrawn at indicated time points were immediately resuspended in 5X SDS sample buffer, boiled at 95 °C for 10 min and then centrifuged. After centrifugation, the clarified supernatant was loaded onto 10 % SDS-PAGE gels. Proteins were then transferred to a nitrocellulose membrane at 100V for 1 hour and probed with polyclonal rabbit anti-CcrM (1:5000 dilution) antibody. Proteins were visualized with goat anti-rabbit (1:5,000) using a Licor detection system (Licor Odyssey CLx). Final images were quantified using imageJ and degradation rates were plotted using Prism.

### Identification of *dcd* suppressors

*ΔdcdΔlon xylX::Pxyl-lon* (CPC624) was grown overnight in PYE supplemented with glucose (0.2%), 100 µL was spread on PYE plates containing xylose (0.2%), incubated at 30°C, and monitored over 4 days. Colonies were picked and re-struck onto agar plates with or without tetracycline to verify Lon integration. Colonies that were still tetracycline resistant were then immunoblotted for Lon and a known Lon substrate ccrM. Clones that had active Lon indicated by low levels of CcrM and high levels of Lon were then grown overnight and then stored at -80°C. Genomic DNA was isolated from these candidates and prepared for Illumina whole genome re-sequencing.

### Illumina sequencing and data analysis

Genomic DNA was isolated using the MasterPure Complete DNA and RNA purification kit according to the manufacturer’s protocol (Epicentre). DNA concentration and quality were assessed using a Qubit Fluorometer (ThermoFisher Scientific). Illumina libraries were generated from isolated genomic DNA using the NexteraXT (Illumina) protocol. Final libraries were multiplexed and sequenced at the University of Massachusetts Amherst Genomics Core Facility on the NextSeq 500 (Illumina). SNPs were detected using breseq, a computational pipeline for identifying mutations relative to a reference using short-read DNA re-sequencing data for microbial-sized genomes (Deatherage and Barrick, 2014).

## Acknowledgements

We thank the Chien, Strieter, Stratton, Serio and Tu lab members for helpful comments and discussion on the manuscript. We thank Lucy Shapiro for providing CcrM antibody. This project was supported by funds from the NIH (R35GM130320 to P.C.). R.Z. was supported in part through the Chemistry Biology Interface Training Program (NIH T32GM08515).

## Author Contributions

Conceptualization, R.Z., P.C.; Investigation, R.Z., H.B.; Writing, R.Z., P.C.; Funding Acquisition, P.C. and B.T.; Supervision, P.C. and B.T.

