## Supplemental Data for "The Lon protease links nucleotide metabolism with proteotoxic stress"

**Supplemental Figure Legends**

**Figure S1. KEGG analysis reveals pathways that are influenced by Lon, Related to Figure 1**

(A) Volcano plot representation of the Tn-seq analysis of *Δlon* vs wt. Negative $Log_{10}$ of the p-value is plotted against $Log_2$ of the fold change in total transposon insertion counts in each gene in wt and *Δlon*. KEGG pathways significantly affected by loss of Lon ($p < 0.05$) are grouped by de-enriched genes (dark blue) or enriched genes (green). Genes within the significant KEGG pathways are colored based FDR <0.05.

Figure S1, Related to figure 1

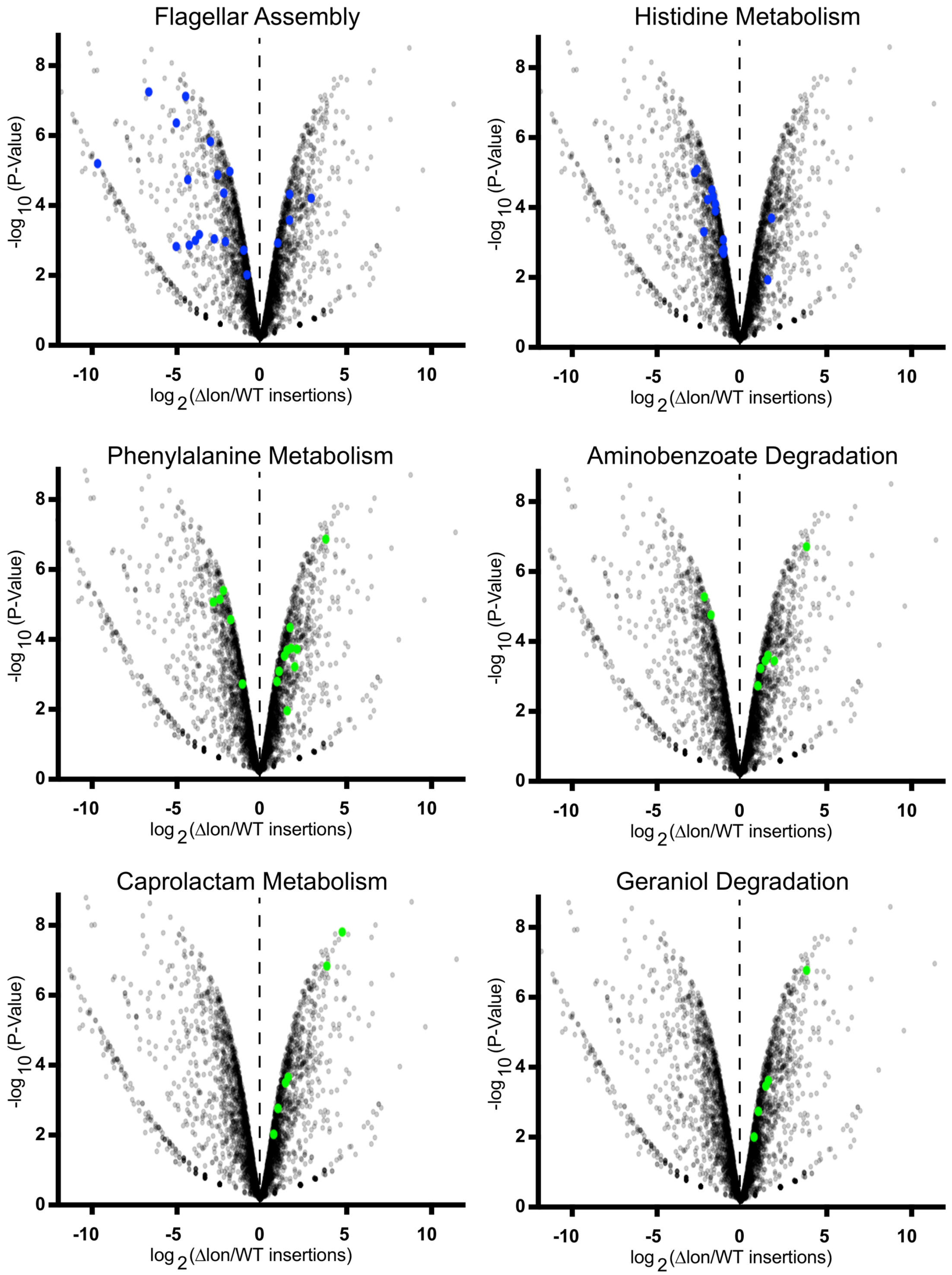

**Figure S2. Deletion of FlhA/FlhB is synthetic lethality with loss of Lon, Related to figure 1**

(A)(B) Plot of transposon insertion frequency and distribution of *flhA* or *flhB* in wildtype (top) and *Δlon* (bottom). For both wildtype and *Δlon* the positive and negative strand insertions are displayed for both biological replicate transposon libraries indicated by a (+) or (-). The 5' insertion sites are displayed on a logarithmic scale.

(E)(F) Clean deletions of *flhA/B* result in reduced motility. Single colonies were inoculated into PYE with 0.3% agar and allowed to grow two days before imaging.

(C)(D) Loss of *flhA/B* in wildtype and *Δlon* confirms synthetic lethality with loss of Lon. Strains were grown overnight and representative images were taken.

Figure S2, Related to figure 1

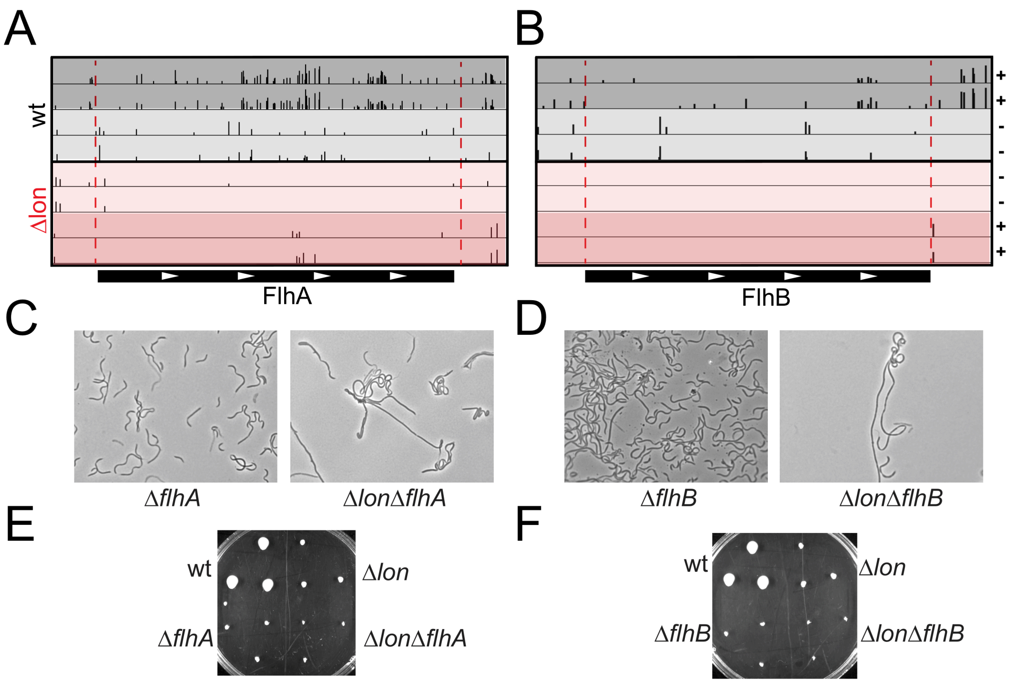

**Figure S3. Predominant suppressors of DCD are loss of function mutations in CTP synthase, Related to figure 3**

(A) Whole genome sequencing analysis of suppressors of *ΔlonΔdcd xylX::Pxyl-lon* strains. Short nucleotide polymorphisms (SNPs) were identified by Breseq analysis and unique mutations are summarized in the data table.

(B) Mutations in CTP synthase are predominant suppressors of DCD deletion. Mutations are clustered in the synthetase domain.

(C) CTP synthase mutations may destabilize protein fold and ATP binding. Model of *Caulobacter* CtpS generated by Phyre2. Suppressor mutations are shown as spheres, insets highlight the local residue and ligand environment of the suppressor mutations.

### Figure S3, Related to figure 3

A

| Gene | Missense Mutation | Annotation |
|---|---|---|
| CCNA_01791 | L72Q (CTG → CAG) | CTP synthase |
| CCNA_01791 | I162S (ATC → AGC) | CTP synthase |
| CCNA_01791 | D245G (GAC → GGC) | CTP synthase |
| CCNA_01791 | V296A (GTC → GCC) | CTP synthase |
| CCNA_03607 | L60P (CTG → CCG) | NrdA |
| CCNA_01966 | L245P (CTG → CCG) | Vitamin B12 dependent RNR |
| CCNA_01966 | I265N (ATC → AAC) | Vitamin B12 dependent RNR |
| CCNA_02087 | N66D (AAC → GAC) | dGTP Triphosphohydrolase |

B

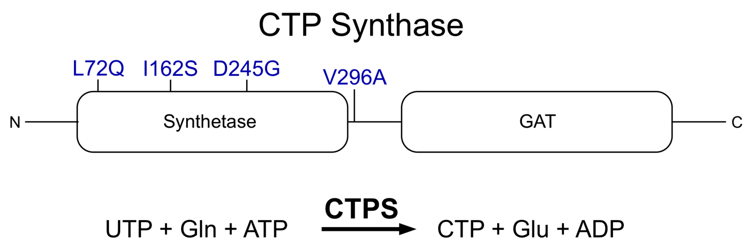


C

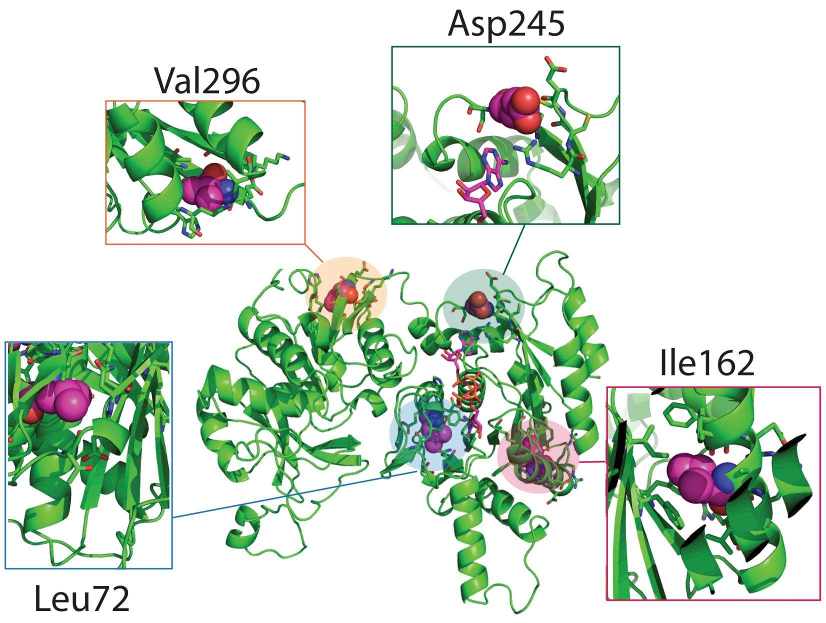

**Figure S4. dUTP/dTTP is sufficient for DCD deletion, Related to Figure 3**

Cell death induced upon Lon induction in a dCTP deaminase deletion can be rescued by addition of 100 µM dTTP/dUTP into solid media but not dCTP. Cells were grown to exponential phase before being serially diluted 10-fold, and spotted onto media supplemented with xylose (0.2%, +Lon) or glucose (0.2%, -Lon) with or without the indicated concentrations of deoxynucleotides.

### Figure S4, Related to figure 3

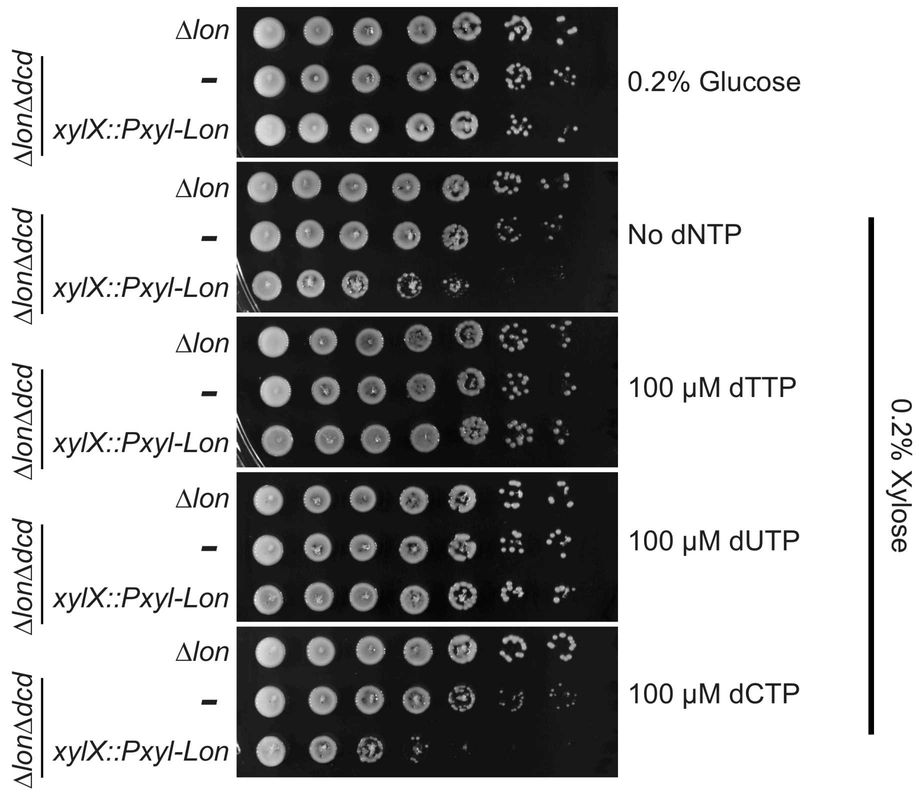

**Figure S5. Lon protease allows for tolerance of DNA damaging agents, Related to Figure 4**

Loss of Lon results in decreased resistance to DNA damage. Strains were grown to exponential phase before being serially diluted 10-folded and spotted onto media supplemented with xylose (0.2%, +Lon), with or without 1 μg/ml mitomycin C (MMC) or 0.00015% methyl methanesulfonate (MMS). Plates were then grown 3 days at 30˚C before imaging.

### Figure S5

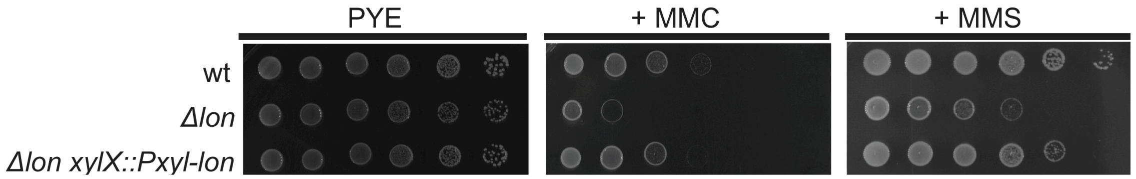

**Figure S6. Uncropped gels and plates used in figure 5.**

**Figure S7. Uncropped western blots used in figure 5.**

### Figure S6, Related to figure 5

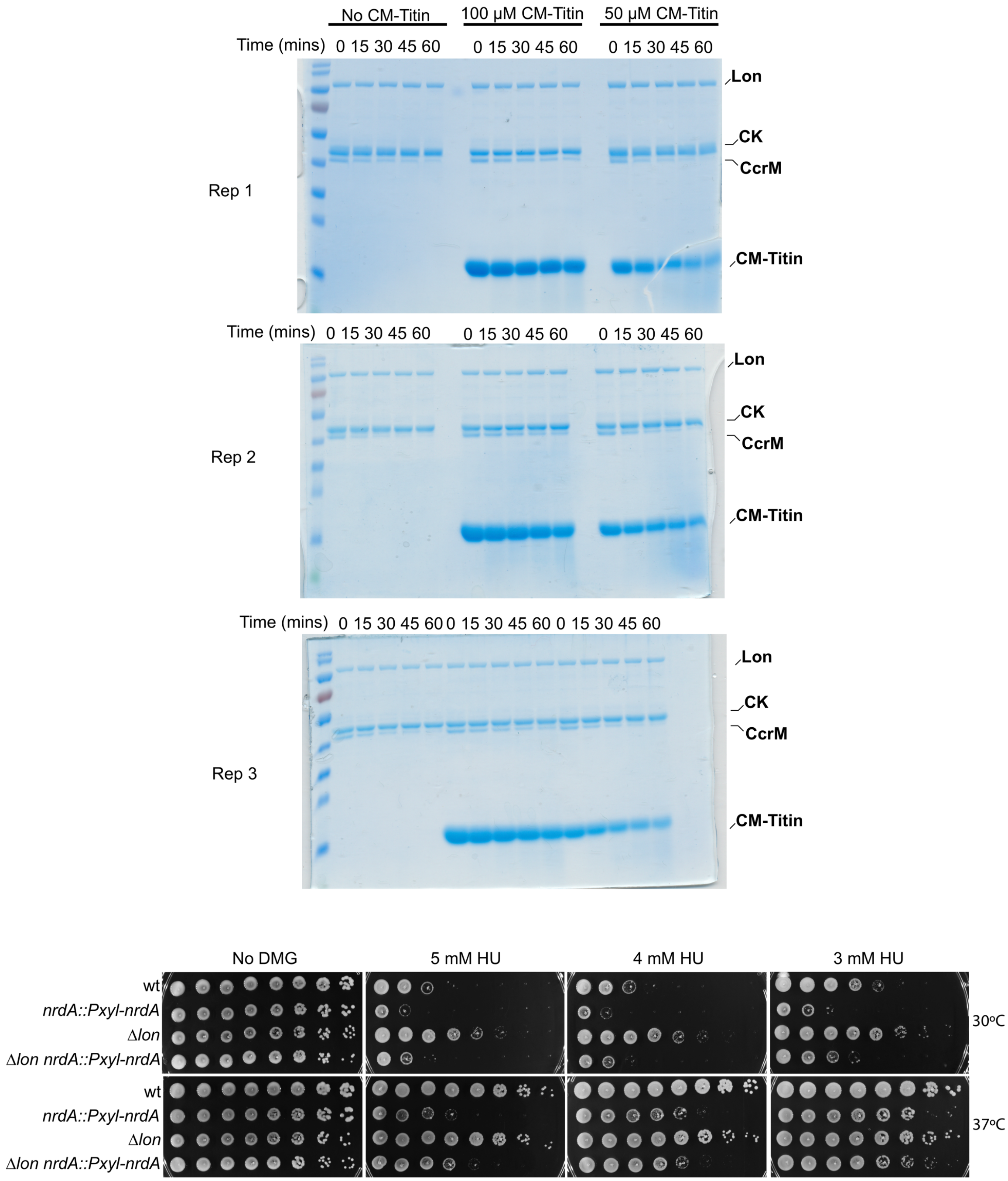

### Figure S7, Related to figure 5

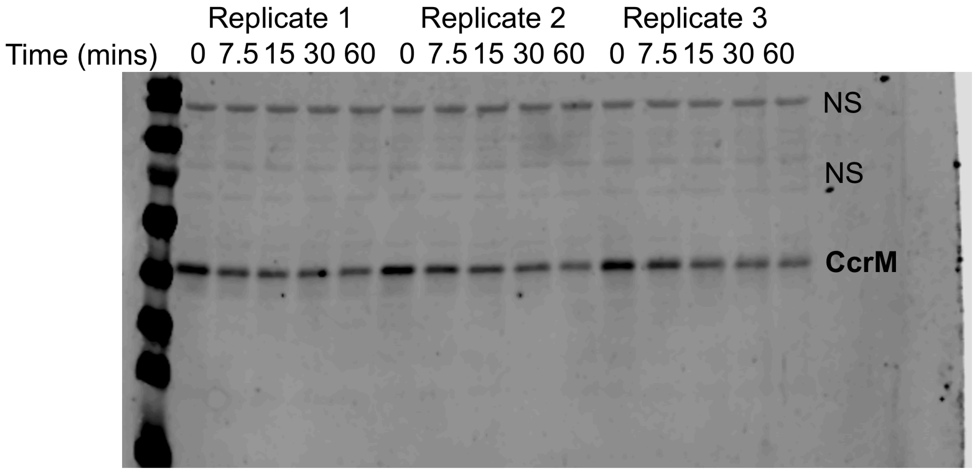


30ºC

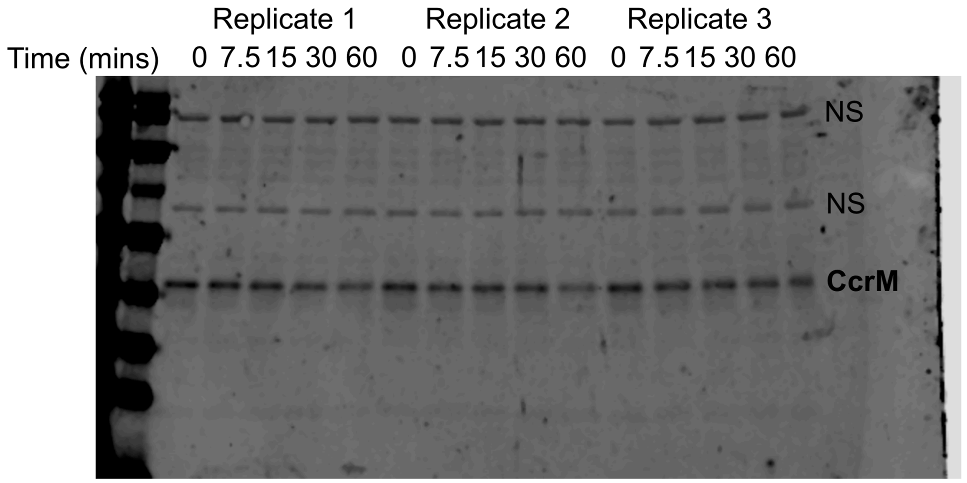


37ºC
